# Identification of a unique TCR repertoire, consistent with a superantigen selection process in Children with Multi-system Inflammatory Syndrome

**DOI:** 10.1101/2020.11.09.372169

**Authors:** Rebecca A Porritt, Lisa Paschold, Magali Noval Rivas, Mary Hongying Cheng, Lael M Yonker, Harsha Chandnani, Merrick Lopez, Donjete Simnica, Christoph Schultheiß, Chintda Santiskulvong, Jennifer Van Eyk, Alessio Fasano, Ivet Bahar, Mascha Binder, Moshe Arditi

## Abstract

Multisystem Inflammatory Syndrome in Children (MIS-C), a hyperinflammatory syndrome associated with SARS-CoV-2 infection, shares many clinical features with toxic shock syndrome, which is triggered by bacterial superantigens. The superantigen specificity for binding different Vβ-chains results in Vβ-skewing, whereby T cells with specific Vβ-chains and diverse antigen specificity are overrepresented in the TCR repertoire. Here, we characterized the TCR repertoire of MIS-C patients and found a profound expansion of TCR Beta Variable gene (TRBV)11-2. Furthermore, TRBV11-2 skewing was remarkably correlated with MIS-C severity and serum cytokine levels. Further analysis of TRBJ gene usage and CDR3 length distribution of MIS-C expanding TRBV11-2 clones revealed extensive junctional diversity, indicating a superantigen-mediated selection process for TRBV expansion. *In silico* modelling indicates that polyacidic residues in TCR Vβ11-2 engage in strong interactions with the superantigen-like motif of SARS-CoV-2 spike glycoprotein. Overall, our data indicate that the immune response in MIS-C is consistent with superantigenic activation.

**Highlights:**

- Multisystem Inflammatory Disease in Children (MIS-C) patients exhibit T cell receptor (TCR) repertoire skewing, with expansion of T cell Receptor Beta Variable gene (TRBV)11-2
- TRBV11-2 skewing correlates with MIS-C severity and cytokine storm
- J gene/CDR3 diversity in MIS-C patients is compatible with a superantigen selection process
- *In silico* modelling indicates TCR Vβ11-2 engages in CDR3-independent interactions with the polybasic insert P_681_RRAR in the SAg-like motif of SARS-CoV-2 spike

## Introduction

Severe acute respiratory syndrome coronavirus 2 (SARS-CoV-2) is the causative agent of coronavirus disease 2019 (COVID-19), which started as an epidemic in China and culminated in a global pandemic. Typical COVID-19-related symptoms include fever, dry cough, breathing difficulties and gastrointestinal (GI) symptoms (Huang et al., 2020). Although most infected adults develop a mild course of the disease, COVID-19 manifests as a severe interstitial pneumonia with hyperinflammation and many extrapulmonary complications in 20% of patients (Cristiani et al., 2020; Tay et al., 2020; Vabret et al., 2020). Severe COVID-19 in adult patients is associated with overactivation of the immune system and increased release of proinflammatory cytokines, a process called “cytokine storm” (Vabret et al., 2020). The percentage of pediatric cases diagnosed with SARS-CoV-2 infection has risen steadily since mid-April from just 2 to 11% of all USA COVID-19 cases (American Academy of Pediatrics, 2020). Children are less likely to suffer from lifethreatening COVID-19 disease, and account for just 3.5% of current COVID-19 hospitalization in the USA (American Academy of Pediatrics, 2020). While rare, children can develop severe disease caused by SARS-CoV-2 infection, presenting with high fever and acute respiratory disease. In addition, beginning in March and April 2020, there have also been reports of children admitted to hospital with a multisystem hyperinflammatory syndrome, presenting with fever, severe abdominal pain, diarrhea, myocardial dysfunction and cardiogenic shock and rash, reminiscent of Kawasaki disease (KD) (Belhadjer et al.; Cheung et al., 2020; Riphagen et al.; Toubiana et al., 2020; Verdoni et al.; Whittaker et al., 2020). The first reports of this new syndrome, now called multisystem inflammatory syndrome in children (MIS-C), were from the UK, and they later emerged from across Europe and the Eastern part of North America (Belhadjer et al.; Cheung et al., 2020; Riphagen et al.; Toubiana et al., 2020; Verdoni et al.; Whittaker et al., 2020). Recently, a similar disease in adults (MIS-A) has also been identified (Morris et al., 2020). Both MIS-C and MIS-A usually require intensive care, however these two entities are not associated with severe respiratory symptoms (Belhadjer et al., 2020; Cheung et al., 2020; Morris et al., 2020; Riphagen et al., 2020; Toubiana et al., 2020; Verdoni et al., 2020; Whittaker et al., 2020). It is interesting to note that approximately a third or fewer of MIS-C patients tested positive for SARS-CoV-2, but the majority had serologic evidence of infection or a history of exposure to COVID-19 (Belhadjer et al.; Cheung et al., 2020; Riphagen et al.; Toubiana et al., 2020; Verdoni et al.; Whittaker et al., 2020). MIS-C may therefore be triggered by an extrapulmonary SARS-CoV-2 infection or be caused by a delayed, post-infectious inflammatory response.

Clinical and laboratory features of MIS-C are remarkably similar to those of Toxic Shock Syndrome (TSS), including severe GI symptoms, neurological symptoms, myocardial involvement, lymphopenia, elevated levels of C-reactive protein (CRP), ferritin, and D-dimers, and increased expression of pro-inflammatory cytokines (Belhadjer et al.; Cook et al., 2020; Low, 2013; Pierce et al., 2020; Riphagen et al.; Verdoni et al.). TSS is driven by bacterial superantigens (SAgs) such as Staphylococcal enterotoxin B (SEB). SAgs are highly potent T cell activators able to engage both T cell receptors (TCRs) and MHC class II (MHCII) molecules (Krakauer, 2019). Typically, bacterial SAgs activate T cells by binding to specific β-chains of TCRs at their variable domain in a complementary-determining region 3 (CDR3)-independent manner. As such, SAgs can bypass the TCR antigen specificity and induce T cell activation and proliferation along with a cytokine storm that results in toxic shock (Krakauer, 2019; Li et al., 1999). SAg binding specificity for different TCR Vβ-chains results in Vβ-skewing, whereby T cells with specific Vβ-chains and diverse antigen specificity are overrepresented in the TCR repertoire of patients upon exposure to SAgs.

Using structure-based computational models, we recently demonstrated that the SARS-CoV-2 spike glycoprotein has a unique insert that displays a SAg-like sequence motif that exhibits a high-affinity for binding TCRs, and may form a ternary complex with MHCII (Cheng et al., 2020). Furthermore, we identified that this SAg-like motif has high sequence and structural similarity to a motif in SEB (Cheng et al., 2020). We further reported TCR Vβ-skewing in adult COVID-19 patients with severe hyperinflammation, consistent with a SAg immune response (Cheng et al., 2020).

Since MIS-C clinical features are quite similar to those of TSS, here we analyzed the TCR repertoire of MIS-C patients to determine if MIS-C is associated with Vβ skewing and junctional diversity as well as signs of SAg activation. Immunosequencing of the peripheral blood samples of MIS-C patients revealed a profound expansion of T cell Receptor Beta Variable gene 11-2 (TRBV11-2), which correlated with MIS-C severity and serum cytokine levels consistent with superantigen triggered immune responses.

## Results

### The TCR repertoires of mild MIS-C patients are richer and more diverse than those of severe MIS-C patients

To determine if MIS-C is associated with TCR repertoire skewing, we collected blood samples from mild MIS-C (n=4), severe MIS-C (n=16) and age-matched febrile control patients (n=15). TCR-sequencing analysis was performed on extracted RNA. Global T cell metrics showed no differences in basic repertoire metrics when comparing all the MIS-C patients (mild and severe) with febrile controls. However, we observed that mild MIS-C patients were characterized by a generally richer TCR repertoire than severe MIS-C patients or febrile controls (**Figure S1A-S1D**). These findings were consistent with adult COVID-19, in which mild cases present with higher TCR richness than age-matched controls indicating diversification of the repertoire in an antigen-dependent manner (Schultheiss et al., 2020).

### Skewing of TRBV genes in MIS-C

We next performed principal component analysis (PCA) of TCR V and J gene usage to determine their global distributions between cohorts. PCA of the TRBV gene repertoires revealed that MIS-C patients clustered apart from febrile controls, whereas the usages of TRAV, TRGV and TRDV genes did not show any skewing (**Figure 1A**). PCA of TRBJ gene usage revealed no differences between febrile controls and MIS-C patients or severity groups (**Figure S2**), indicating that a selective pressure was exclusively exerted on the V gene distribution.

**Figure 1.**
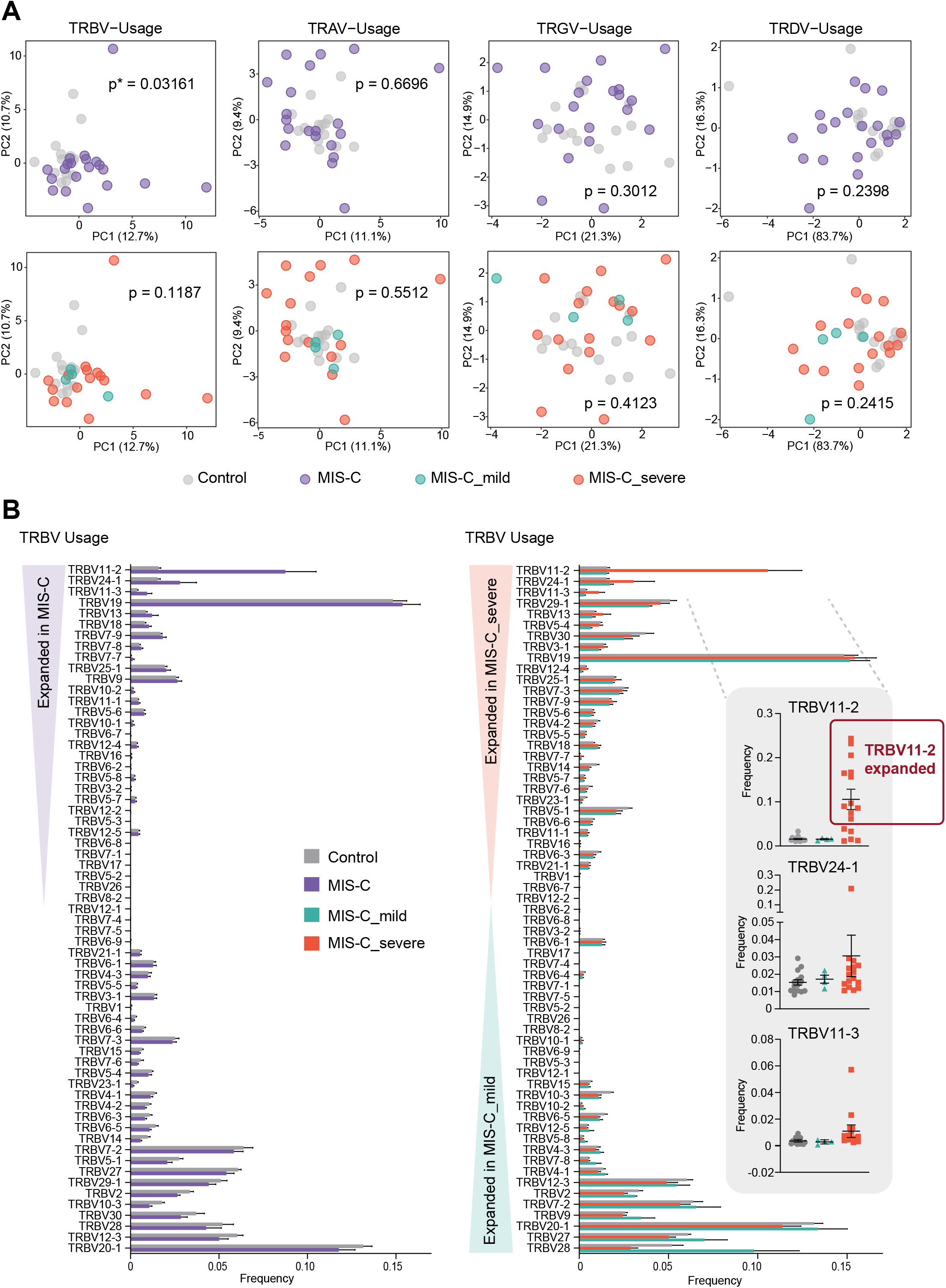
TRBV Driven Repertoire Skewing in MIS-C Patients. **(A)** Principal component analysis (PCA) of differential TRBV, TRAV, TRGV and TRDV gene usage in MIS-C patients (n=20) versus febrile controls (n=15), and between mild MIS-C (n=4) and severe MIS-C (n=16) patients. Statistical analysis: Pillai–Bartlett test of MANOVA of all principal components. **(B)** Frequencies of differentially used TRBV genes in MIS-C and febrile control patients. Bars indicate mean ± SEM. Individuals considered as TRBV11-2 expanded are marked with a red box.

### Severe MIS-C TCR repertories are characterized by a profound expansion of TRBV11-2

Since MIS-C patients exhibit strong TRBV skewing, we next performed differential gene usage analysis between mild MIS-C, severe MIS-C and febrile controls. Several TRBV genes were overrepresented in the overall MIS-C patient group. Specifically, TRBV11-2, TRBV24-1 and TRBV11-3 were enriched in MIS-C patients relative to febrile control patients (**Figure 1B**). Further comparisons between subgroups showed that this enrichment was restricted to individuals with severe disease. In contrast, TRBV-28 was exclusively enriched in mild MIS-C patients. We previously found that TRBV24-1 was overrepresented in a cohort of adult COVID-19 patients with severe, hyperinflammatory courses compared to patients with mild COVID-19 (Cheng et al., 2020). Figure 2A shows the distribution of TRBV11-2, TRBV24-1 and TRBV11-3 genes on an individual repertoire level. This analysis shows that expansion of the TRBV11-2 compartment predominantly occurs in individuals with severe disease. Furthermore, analysis of serum cytokines from matching samples (18 of 20 MIS-C patients) demonstrated significant correlation between TRBV11-2 usage and TNF-α, IFN-γ, IL-6 and IL-10 levels (**Figure 2B**), indicating that TRBV11-2 expansion is associated with the cytokine storm.

**Figure 2.**
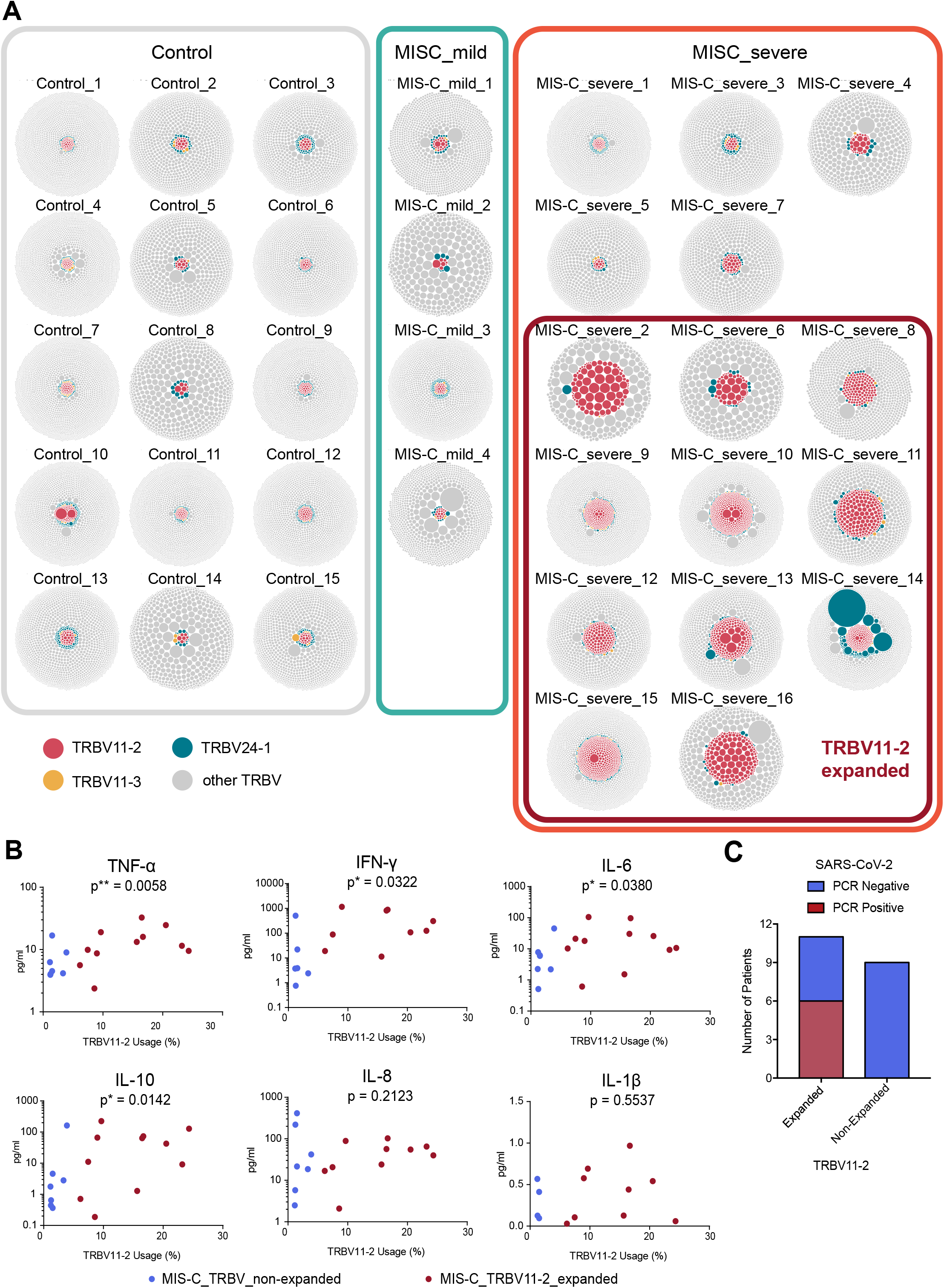
Clonal Distribution in Individual Repertoires of MIS-C Patients. **(A)** Visualization of clonal richness and distribution in individual productive repertoires from mild (n=4) and severe (n=16) MIS-C patients and febrile controls (n=15). One bubble represents one amino acid clonotype, and the area size of the bubble indicates its fraction within the repertoire. Clonotypes using TRBV11-2, TRBV24-1 or TRBV11-3 are highlighted. Individuals considered as TRBV11-2 expanded are marked with a red box. **(B)** Correlation of serum cytokine levels with TRBV11-2 usage. Statistical analysis: Spearman r correlation test. **(C)** Distribution of nasopharyngeal SARS-CoV-2 PCR results amongst patients with or without TRBV11-2 expansion.

TRBV24-1 and TRBV11-3 expansions were less pronounced and restricted to fewer individuals. One patient, “MIS-C_severe_14” showed massive expansion of TRBV24-1 (**Figure 2A**). This patient was 15 years old, which might explain why their TCR repertoire was closer to that previously observed in adult patients. Overall, children between seven and 16 years of age showed strong TRBV11-2 expansions associated with the severe course of the disease (**Figure S3**). However, no age drifts in TRBV11-2, TRBV24-1 or TRBV11-3 distribution were found in the TCR repertoires of 254 individuals, indicating that age does not underlie TRBV11-2 skewing in severe MIS-C patients (**Figure S4**).

Of the MIS-C patients used in this study, 95% were serology positive for SARS-CoV-2, while only 30% were PCR positive by nasopharyngeal swab. Interestingly, however, PCR positivity seemed to correlate with TRBV11-2 expansion. We found that 55% of patients with TRBV11-2 expansion were PCR positive for SARS-CoV-2, whereas none of the patients without expansion were PCR positive for SARS-CoV-2 (**Figure 2C**). These data suggest an association of TRBV11-2 expansion with active SARS-CoV-2 infection.

### Junctional diversity in MIS-C patients with expanded TRBV11-2

SAg interactions involve the V gene, but spare the CDR3 of the TCR. If the TRBV11-2 expansions observed in patients with severe MIS-C were due to SARS-CoV-2 acting as a SAg, the diversity of the V(D)J junction in TCR with TRBV11-2 usage should be high. To investigate junctional diversity, we first searched for CDR3 overlaps between different MIS-C patients. This analysis showed no overlaps between patients with TRBV11-2 expansions, suggesting high CDR3 diversity in this cohort (**Figure 3A**). Repetitive blood samples taken from the same healthy donor were used as control, showing overlapping repertoires at the different sampling time points. In our global analysis, TRBV J gene usage was not biased in MIS-C patients (**Figure S2B**).

**Figure 3.**
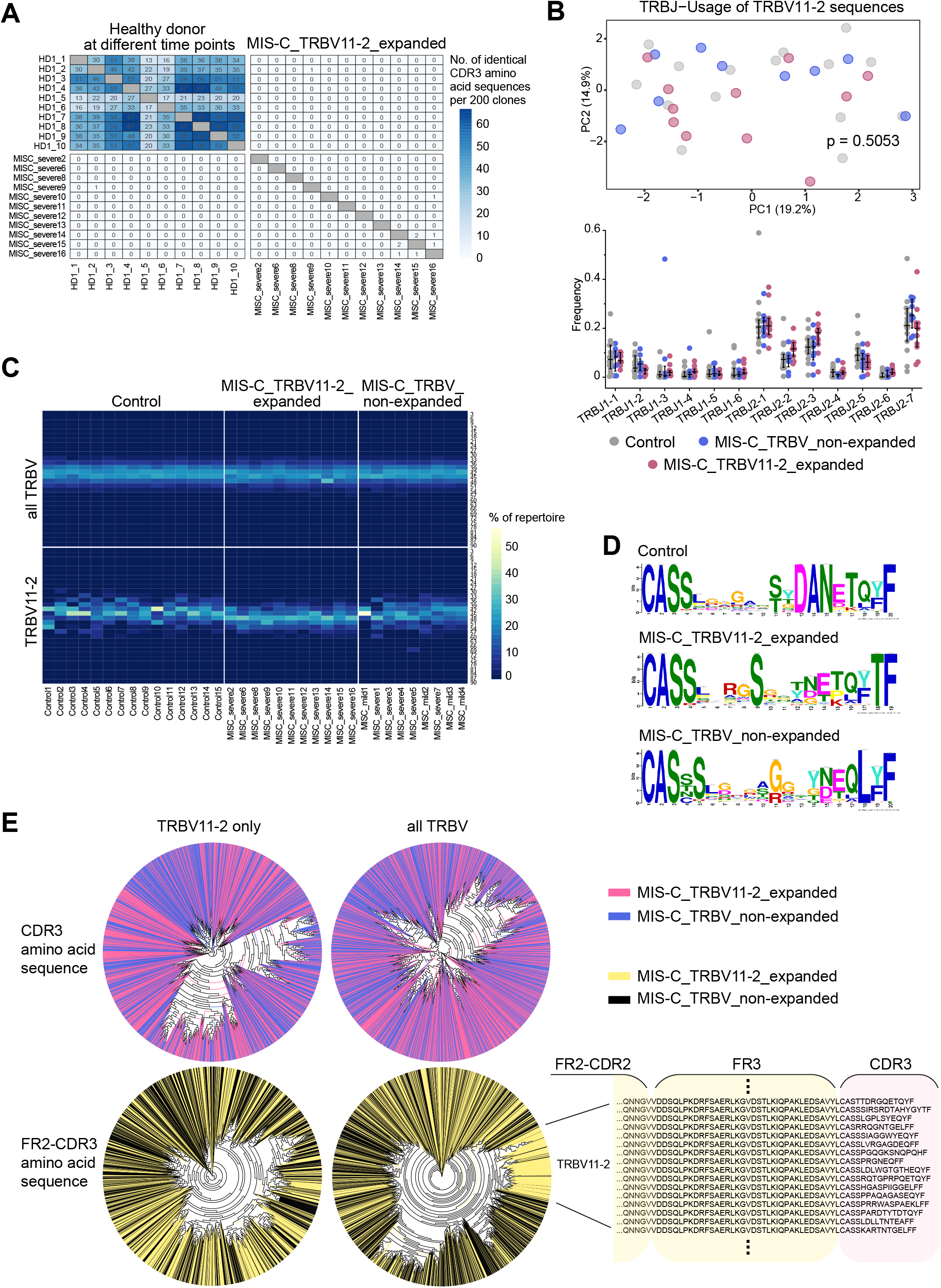
Junctional Diversity in MIS-C Patients with TRBV11-2 Expansions. **(A)** Overlap of CDR3 amino acid clonotypes per 200 sequences in repetitive samples of a healthy individual and between repertoires of MIS-C patients with TRBV11-2 expansion. **(B)** PCA and differential usage of J genes rearranged with TRBV11-2 in MIS-C patients and febrile controls. Bars indicate mean ± SD. Statistical analysis: Pillai–Bartlett test of MANOVA of all principal components. **(C)** Heat map of productive CDR3 length distribution in the repertoires of MIS-C patients (n=20) and febrile controls (n=15). **(D)** CDR3 diversity in MIS-C patients with or without expanded TRBV11-2 and febrile controls displayed as positional weight matrix generated using GLAM2. **(E)** Unsupervised phylogenetic analysis of the amino acid sequences spanning FR2 to CDR3 versus CDR3 alone in the top 100 clones of MIS-C repertoires (n=20), either comprising the complete TRBV sequence pool or TRBV11-2 sequences only.

To further investigate junctional diversity specifically for the V genes expanded in MIS-C cases, we extracted all the J genes rearranged with TRBV11-2 from the repertoires of severe and mild MIS-C patients, and compared these to J genes extracted from age-matched febrile controls. TRBJ2-1,2-2, 2-3 and 2-7 were the most frequent combinations for TRBV11-2, however, we did not see a difference in TRBJ-usage between the different subcohorts, suggesting a full diversity of J genes rearranged with TRBV11-2 (**Figure 3B**). Moreover, we found an even distribution of CDR3 lengths within the fraction of TCRs using TRBV11-2, suggesting no CDR3-driven clonal selection (**Figure 3C**). CDR3 diversity in TCRs using TRBV11-2 was also confirmed by Gapped Local Alignment of Motifs (GLAM) analysis (**Figure 3D**) as well as by unsupervised analysis of phylogenetic trees constructed from the top 100 clones of each repertoire (**Figure 3E**). The latter analysis showed clustering of expanded TRBV11-2 sequences only if the full sequence from framework 2 (FR2) to CDR3 was used for construction, but not if the clustering analysis was limited to the CDR3 sequence (**Figure 3E**). Together, our results suggest that patients with MIS-C show expansion of TCRs using distinct V genes, along with J gene/CDR3 diversity in these rearrangements, compatible with a SAg selection process.

### TRAV8-4 is overrepresented in patients with expanded TRBV11-2

While no global skewing was observed for TRA, TRG or TRD V or J gene distributions, we reasoned that minor skewings, especially of the TRAV repertoire that is associated with TRBV, may have been missed due to the global PCA approach. Since some SAgs can interact with both the TCR Vβ and Vα chains (Saline et al., 2010), we searched for skewing in TRAV, TRGV or TRDV genes associated with TRBV11-2 by investigating their distributions in MIS-C patients with and without TRBV11-2 expansions. While TRGV and TRDV genes did not show any skewed distribution, there was a significant TRAV skewing in MIS-C patients with and without TRBV11-2 expansions (**Figure 4A**). Further analyzes suggested that no single TRAV gene was associated with expanded TRBV11-2, potentially reflecting the lack of skewing in mild versus severe MIS-C cases, but a number of different TRAVs were overrepresented in MIS-C patients with TRBV11-2 expansions and a severe clinical course (**Figure 4B**). TRAV8-4 was the most expanded gene, while expansions of TRAV17 and TRAV22 were less pronounced. These data suggest that the SAg interaction also involves the TRAV genes in MIS-C patients.

**Figure 4.**
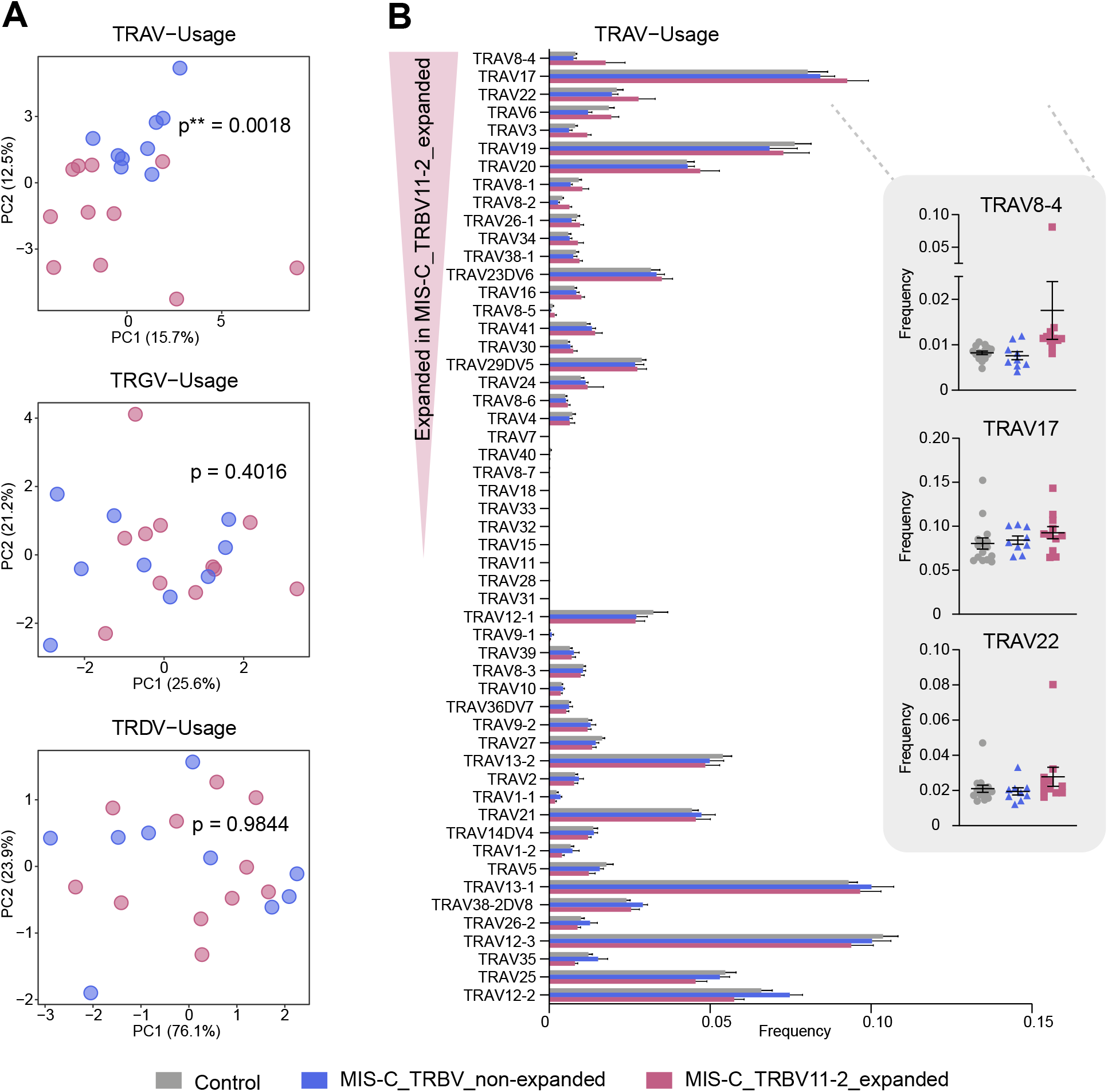
TRAV Repertoire Skewing and Differential Gene Usage in MIS-C Patients. **(A)** PCA of differential productive TRAV gene usage in MIS-C patients (n=20) versus febrile controls (n=15). MIS-C subgroups: mild MIS-C (n=4), severe MIS-C (n=16), MIS-C without TRBV11-2 shift (n=9), MIS-C with TRBV11-2 shift (n=11). Statistical analysis: Pillai–Bartlett test of MANOVA of all principal components. **(B)** Frequencies of differentially used TRAV genes in MIS-C with/without TRBV11-2 shift as compared to febrile controls.

### *In silico* modelling indicates TCR Vβ11-2 engages in a CDR3-independent interaction with the SAg-like motif of SARS-CoV-2 spike

We previously demonstrated though *in silico* modeling the presence of a SAg-like motif on the SARS-CoV-2 spike protein, distinguished by its high sequence- and structural-similarity to a segment of SEB (Cheng et al., 2020). Furthermore, Vβ24-1 was shown to have a high affinity for binding to the SAg-like motif. Given the extensive expansion of TRBV11-2 in MIS-C patients, we examined *in silico* whether the Vβ11-2 chain could engage in a strong interaction with the SAg-like motif on spike. We first examined whether the SARS-CoV-2 spike glycoprotein could bind to Vβ11-2. To this aim, we used a TCR structure resolved in a crystal structure of the monoclonal T cell line DD31 TCR complexed with HLA B*0801 (Nivarthi et al., 2014), the β-chain variable domain of which is sequentially identical to the product Vβ11-2 of TRBV11-2. Following our previous approach (Cheng et al., 2020), we generated a series of structural models for the SARS-CoV-2 spike–TCR complex using the docking software ClusPro (Kozakov et al., 2017), and analyzed the models grouped in structurally similar clusters to detect recurrent binding patterns (**Figure 5**). The analysis led to two hot spots that bound Vβ11-2 on the spike protein: (i) within the SAg-like motif (residues Q677 to R685) and (ii) near the neurotoxin-like motif (residues E340 to R357) identified earlier (Cheng et al., 2020). Interestingly, both hot spots contained polybasic residues at their binding epitopes, P_681_**RR**A**R** and N_355_**RKR**I respectively. Notably, the amino acids D67 and D68 near the CDR2 of Vβ11-2 played a significant role in binding to these two motifs on the spike, thus serving as a paratope in more than 90% of the generated models for the TCR-spike complex. While we do not exclude the potential role of other toxin-like motifs in stimulating hyperinflammatory responses, we focused on the SAg-like motif Q677-R685 reported in our previous study, whose deletion has been shown to attenuate the severity of the disease (Lau et al., 2020).

**Figure 5:**
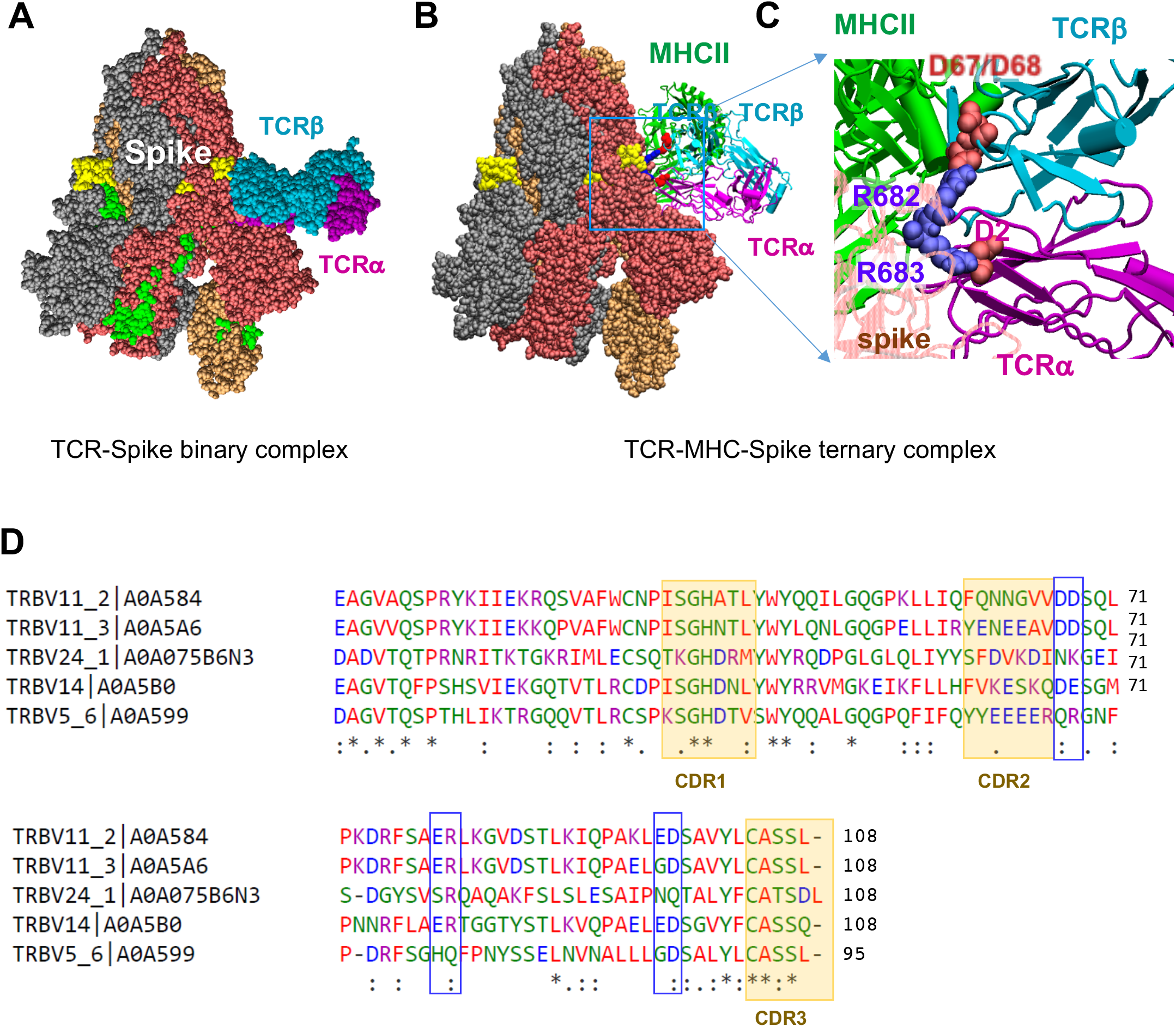
Complex Formation Between SARS-CoV-2 spike, TCR Containing Vβ11-2, and MHCII, and Comparative Analysis of TCR Vβ Sequences Homologous to Vβ11 −2. **(A)** Binding of TCR (with Vβ chain sequentially identical to that of TRBV11-2 gene product) to the SAg-like region of SARS-CoV-2 spike. The TCR α- and β-chains are shown in *magenta* and *cyan,* respectively. The β-chain tightly binds the SAg-like region (E661 to R685; colored *yellow).* The spike subunits are colored *dark red, beige,* and *gray;* and the neurotoxin motif (299-356), *green.* (**B-C**) Ternary complex between spike, the same TCR and MHCII (*green* and the close-up view of the interfacial interactions between two basic residues, R682 and R683, on the SAg-like region of spike and the acidic residues (D67 and D68) of the TCR Vβ (D67 and D68) and TCRα. (**D**) Multiple sequence alignment (MSA) of the TCR Vβ chains distinguished by Vβ-skewing in MIS-C (TRBV11-2, TRBV11-3 and TRBV24-1) and severe adult COVID-19 patients (TRBV24-1, TRBV14 and TRBV5-6). The MSA was generated by Clustal Omega (Sievers et al., 2011). CDR1, CDR2 and CDR3 are indicated by *orange shades.* The regions in *blue boxes* indicate the highly conserved paratopes in TRBV11-2 and TRBV14. See more extended MSA in Figure S5.

Figure 5A illustrates the binding of the TCR containing Vβ11-2 domain to the SARS-CoV-2 spike at the SAg-like region containing the polybasic insertion, P_681_**RR**A**R**, unique to SARS-CoV-2 among all SARS family coronaviruses. Notably, the complex is stabilized by CDR3-independent interactions. Instead D67-D68 in Vβ11-2 (**Figure 5B, C**) and Vα chain D2 tightly bind the polybasic insert of the spike via salt bridges between these oppositely charged pairs of residues. Modeling and simulations for Vβ14 (encoded by a skewed TRBV gene, TRBV14, identified in adults of severe Covid-19 patients (Cheng et al., 2020); not shown) also revealed the involvement of the corresponding acidic residues D67 and E68 in binding the same epitope. We also observed that D97-D98 in both Vβ11-2 and Vβ14 were also able to bind to the PRRAR region, albeit but with lower probability than D67-D68. Further simulations showed that the same binding site simultaneously accommodates the binding of a MHCII molecule with minimal rearrangement in the binding pose of the TCR (**Figure 5B**), suggesting that a ternary interaction is possible between the TCR, MHCII and spike, provided that there is no steric overlap between the MHCII and the T cell membrane to which the TCR is bound.

To further assess the specificity of the polyacidic residues near the CDR2, including D67 and D68 in Vβ11-2, or D67 and E68 in Vβ14, we searched the database of TRBV genes compiled in the IMGT Repertoire (Lefranc et al., 2009). We identified a total of ten human TRBV genes encoding polyacidic residues near CDR2: TRBV7-3, TRBV11-1, TRBV11-2, TRBV11-3, TRBV12-3, TRBV12-4, TRBV12-5, TRBV14, TRBV16, and TRBV18. The corresponding multiple sequence alignment is presented in Figure S5. With the exception of TRBV11-1, 11-2, and 11-3, all others have been identified as specific Vβ chains specific to SAgs secreted by *Staphylococcus aureus* and other select *Staphylococcal* species that lead to TRBV skewing in TSS patients (Tuffs et al., 2018). Notably, TRBV11-2 and TRBV14 share three homologous binding paratopes *(blue boxes* in **Figure 5D** and **Figure S5**) and may share similar mechanisms of action.

Finally, while the amino acid sequences corresponding to TRBV11-1, −2, and −3 are highly similar all being TRBV11 subtypes, their most important differences reside in their CDR2 sequence. Notably, TRBV11-2 CDR2 encodes a series of hydrophobic and polar residues that would favor interfacial contacts with the spike which would bury the hydrophobic residues; whereas the CDR2s of TRBV11-1 and 11-3 each contain two or three acidic residues in addition to D67-D68. Such a cluster of 4-5 negatively charged residues that are closely neighboring in sequence (and space) may give rise to destabilizing effects at possible protein-protein interfaces, and would favor instead an exposure to the solution. Thus, the sequence difference between these TRBV11 subtypes seems to explain the significantly higher binding propensity of TRBV11-2, compared to TRBV11-1 and −3.

## Discussion

Here, we report the identification of a SAg-induced TCR signature in MIS-C patients. A similar finding was also previously observed in adults with severe hyperinflammatory COVID-19 (Cheng et al., 2020). COVID-19 and MIS-C are both hyperinflammatory syndromes associated with SARS-CoV-2 infection, and we have reported the presence of a SAg motif in SARS-CoV-2 spike protein (Cheng et al., 2020). In MIS-C patients, TRBV11-2 was profoundly expanded and correlated with disease severity, whereas TRBV24-1 and TRBV11-3 were overrepresented to a much lesser degree. The TRBV11-2 expansions observed in many of the MIS-C patients with severe clinical courses followed the classical SAg expansion pattern, since they do not rely on the VDJ junction. Furthermore, the structure-based *in silico* modeling indicated that TCR Vβ11-2 can engage in a CDR3-independent interaction with the SAg-like motif of in SARS-CoV-2 spike protein. These data suggest that SARS-CoV-2 spike may act as a superantigen to trigger the development of MIS-C as well as cytokine storm in adult COVID-19 patients, with important implications for the development of therapeutic approaches.

MIS-C patients present with lymphopenia and hallmarks of hyperinflammation, including elevated levels of CRP, ferritin, troponin, and D-dimers and increased expression of pro-inflammatory cytokines IL-6, IL-1β, IFN-γ and IL-8 (Belhadjer et al.; Carter et al., 2020; Cheung et al., 2020; Consiglio et al., 2020; Pierce et al., 2020; Riphagen et al.; Toubiana et al., 2020; Verdoni et al.; Whittaker et al., 2020). We observed a significant correlation of IFN-γ, TNF-α, IL-6 and IL-10 with TRBV11-2 usage, indicating TRBV11-2 expansion may be driving the cytokine storm. Studies examining the T cell compartment in MIS-C are limited, however T cell frequencies appear to be reduced relative to healthy controls and pediatric COVID-19 patients (Carter et al., 2020; Consiglio et al., 2020; Gruber et al., 2020; Vella et al., 2020). One study employing mass cytometry (CyTOF) immunophenotyping found that the distribution of naïve, central memory, effector and effector memory T cells expressing CD45RA (TEMRA) subsets in MIS-C patients were similar to those of healthy controls (Gruber et al., 2020). However, another study using flow cytometry immunophenotyping found that MIS-C patients had increased effector memory subsets as a fraction of total CD4^+^ or CD8^+^ T cells compared with controls (Consiglio et al., 2020). Furthermore, they found that both non-MIS-C and MIS-C SARS-CoV-2 infected children had lower frequencies of follicular helper T cells (Tfh cells) compared with controls (Consiglio et al., 2020). Interestingly, the bacterial superantigen SEB can induce effector memory CD4^+^ and CD8^+^ T cell expansion but has little effect on Tfh cells (Rudolph et al., 2018). Another study found that the T cell compartment of MIS-C was highly activated, particularly a subset of vascular patrolling T cells (CX3CR1^+^ T cells) (Vella et al., 2020). Furthermore, they found an increased frequency of CD8^+^ T cells co-expressing PD-1 and CD39, suggesting prolonged T cell stimulation (Vella et al., 2020).

While studies have shown that MIS-C patient plasma can neutralize SARS-CoV-2 infection (Gruber et al., 2020), children with either COVID-19 or MIS-C were found to have reduced neutralizing activity compared with adult COVID-19 patients (Weisberg et al., 2020). MIS-C patients have high levels of IgG and IgA, and low IgM, indicating class switching (Gruber et al., 2020). Immune profiling of MIS-C patients has also identified the presence of IgG and IgA autoantibodies that recognize immune-cell, endothelial cell, myocardial and GI autoantigens, indicating a breakdown of B cell tolerance (Consiglio et al., 2020; Gruber et al., 2020). These findings are in line with the currently proposed model of a SAg response driving MIS-C, as the cytokine storm induced by SAg-stimulated T cells may act to enhance autoreactive B cell responses. Furthermore, SAgs can mediate T cell interactions with B cells by binding to B cell MHCII, leading to their proliferation and differentiation into Ig secretory cells (Mourad et al., 1989). These direct interactions can promote both polyclonal and antigen-specific B cell activation (He et al., 1992; Tumang et al., 1991). Endothelial cells (ECs) also express MHCII and ICAM-1 in response to IFNγ stimulation, and mediate SAg-induced proliferation and activation of T cells in a similar manner to antigen presenting cells (Krakauer, 1994). Furthermore, SAg-activated cytotoxic T cells can mediate killing of inflammatory ECs (Riesbeck et al., 1998). The release of autoantigens by ECs lysis and cytokine storm-induced tissue damage, in addition to increased B cell responses, likely contributes to autoantibody production. However, whether these autoantibodies participate in MIS-C pathogenesis or are a byproduct of SAg-induced cytokine storm and tissue damage remains to be determined.

Approximately a third or fewer of MIS-C patients were reported to test positive for SARS-CoV-2 by nasopharyngeal PCR (Belhadjer et al.; Cheung et al., 2020; Riphagen et al.; Toubiana et al., 2020; Verdoni et al.; Whittaker et al., 2020). This may suggest that the SARS-CoV-2 SAg causes a post-infectious, delayed hyperinflammation response in certain children. It is also possible that, despite a negative nasopharyngeal PCR test, the virus may still be present in the GI tract. MIS-C patients demonstrate unusually severe GI symptoms, abdominal pain, vomiting and diarrhea (Belhadjer et al., 2020; Riphagen et al., 2020; Verdoni et al., 2020; Whittaker et al., 2020) and it is now known that SARS-CoV-2 can infect the GI tract with evidence for persistent fecal shedding of the virus in children despite testing negative by nasopharyngeal swab (Xu et al., 2020). Interestingly, in our cohort, TRBV11-2 expansion was associated with PCR positivity, suggesting a direct triggering of T cell activation by the virus. However, one third of severe MIS-C patients had neither T cell expansion nor active infection. It is possible that in those patients, the disease has progressed further such that SAg-activated T cells have been depleted due to activation-induced cell death, and a post-infectious immune or autoimmune response is occurring. Supporting this possibility, SAgs have been implicated in various autoimmune diseases (Hurst et al., 2018). For example, the SAg-producing bacteria, *Streptococcus pyogenes,* can induce post-infectious acute rheumatic fever two to four weeks after pharyngeal infection (Hurst et al., 2018). One hypothesis is that SAgs induce autoimmunity by triggering self-reactive T cells (Hurst et al., 2018; Li et al., 1999). As discussed above, SAg activation of autoantibodies in MIS-C may also contribute to a post-infectious autoimmune response.

Severe COVID-19 patients also present with lymphopenia, hyperinflammation and multiorgan involvement (Tay et al., 2020). Similar to adult COVID-19, we found mild cases of MIS-C present with higher TCR repertoire richness and diversity than age-matched controls (Schultheiss et al., 2020). Our previous studies on adult patients also found an expansion of TRBV genes consistent with SAg activation, including TRBV24-1 (Cheng et al., 2020). While TRBV24-1 overrepresentation was common in severe COVID-19 and MIS-C, TRBV11-2 overrepresentation was unique to MIS-C. TCR diversity is lost with increasing age (Simnica et al., 2019a), however we found no correlation of TRBV11-2 or TRBV24-1 usage with age in healthy controls, which suggests this may not be a mechanism driving this difference. Interestingly, SEB has differential effects on human peripheral T cells depending on the donor age, with enhanced activation occurring in adult compared to pediatric donor cells (Rudolph et al., 2018). Whether intrinsic differences in T cells lead to altered TRBV expansion in children and adults in response to SARS-CoV-2 remains to be determined.

While early studies showed that bacterial SAgs activate T cells by binding to the β chain of the TCR (Choi et al., 1989; Fraser and Proft, 2008; Scherer et al., 1993), more recent studies revealed that they can bind to either α-or β-chains, or both (Saline et al., 2010). We found an overrepresentation of TRAV8-4 in patients with TRBV11-2 expansion, which indicates the Vα chain of the TCR may help to stabilize the interaction of spike with Vβ11-2. Indeed, the *in silico* models constructed for TCR-spike complexes, utilizing Vβ11-2 or skewed TRBV genes identified in adults of severe Covid-19 patients (Cheng et al., 2020), also indicated interactions between the SAg-like region and the Vα chain (see the salt bridge formation between D2 and R683 in **Figure 5C**). However, the strongest interactions take place here with the CDR2 of Vβ11-2.

MIS-C was first described as a “Kawasaki-like” disease due to some overlapping clinical symptoms. However, it soon became evident that MIS-C is distinct from both KD and KD-shock based on demographics, clinical and laboratory parameters, and immune profiles (Consiglio et al., 2020; Whittaker et al., 2020). Notably, it was previously proposed that the triggering agent of KD might be a SAg. Indeed, several studies have reported that, compared with healthy control individuals, patients with KD exhibit a skewed Vβ T cell repertoire and increased frequencies of circulating Vβ2 and Vβ8-1 T cells (Abe et al., 1992; Abe et al., 1993; Curtis et al., 1995) and Vβ2 in the mucosa of the small intestine (Yamashiro et al., 1996). However, these results could not be reproduced, and later studies did not confirm TCR skewing and expansion of the described TCR clones (Mancia et al., 1998; Pietra et al., 1994; Sakaguchi et al., 1995). More recent work from multiple teams has shown that B cells as well as CD8^+^ T cells are involved in KD pathogenesis (Brown et al., 2001; Choi et al., 1997; Rowley et al., 2005; Rowley et al., 2000; Rowley et al., 2001), leading to the more widely accepted theory that KD is more likely a conventional antigen-driven response that can be triggered by multiple conventional antigens.

MIS-C patients have responded well to treatment with IVIG and steroids (Belhadjer et al.; Riphagen et al.; Verdoni et al.). Importantly, IVIG is also used for treatment of TSS and is known to contain anti-SEB antibodies (Nishi et al., 1997), and presumably antibodies against other SAgs. Cross reaction of SAg antibodies has been observed, implying shared epitopes (Bavari et al., 1999). The success of IVIG in MIS-C patients may therefore be due to cross-neutralization of the spike by SAg-specific antibodies. Approximately 80% of individuals over age 12 harbor anti-SEB antibodies, however protective SEB titers fall in older adults after age 70 (LeClaire and Bavari, 2001; McGann et al., 1971). These metrics may explain why the elderly are less protected against severe COVID-19. As the spike protein of SARS-CoV-2 contains a SAg-like motif similar to SEB (Cheng et al., 2020), this also raises the possibility of using humanized monoclonal anti-SEB Abs (Dutta et al., 2015; Larkin et al., 2010) for the treatment of MIS-C, and designing peptide mimetics to block SAg interactions, as has been shown for SEB (Arad et al., 2011).

Although we have described TCR skewing in severe adult hyperinflammatory COVID-19 patients (Cheng et al., 2020) and now in children with MIS-C, consistent with immune responses against SAg structures, direct superantigenic activity of the SARS-CoV-2 spike protein awaits additional viral tools. SARS-CoV-2 spike protein is a membrane bound and heavily glycosylated trimer which undergoes multiple post-translational modifications and conformational changes (Wrapp et al., 2020). Expression of a stable recombinant spike protein requires mutation of R_682_RAR_685_ (Wrapp et al., 2020), which removes the putative SAg-like structure. Furthermore, exposure of SARS-CoV-2 SAg motif depends on certain conformations of the spike protein, which are difficult to replicate in an *in vitro* system. However, a number of very recent studies implicate the P_681_RRAR_685_ SAg motif we identified (Cheng et al., 2020) in viral pathogenesis (Cantuti-Castelvetri et al., 2020; Daly et al., 2020; Lau et al., 2020). SARS-CoV-2 S1/S2 cleavage exposes the P_681_RRAR_685_ motif at the C-terminus of the S1 trimer. The same polybasic site has been recently shown to bind to the cell surface receptor Neuropilin-1 (NRP1) (Daly et al., 2020). Remarkably, NRP1 binding is important for SARS-CoV-2 cellular infectivity, as blockade of this interaction by RNAi or mAb against NRP1 significantly reduced *in vitro* SARS-CoV-2 cellular entry (Cantuti-Castelvetri et al., 2020; Daly et al., 2020). Moreover, in an independent study, a SARS-CoV-2 variant with a natural deletion of the S1/S2 furin cleavage site, which contains the P681RRAR685 motif, resulted in attenuated viral pathogenesis in hamster models (Lau et al., 2020). Therefore, targeting this SAg motif might provide a new therapeutic avenue to treat and prevent severe COVID-19, MIS-C and MIS-A.

Overall, our data show a unique TCR repertoire in MIS-C patients, characterized by a profound expansion of TRBV11-2 which is consistent with SAg-induced T cell skewing, and provides a functional relevance to the recently identified SAg-like motif in SARS-CoV-2 (Cheng et al., 2020). Future investigations aiming to characterize the phenotype and functional properties of T cells utilizing TRBV11-2 in MIS-C patients are needed to provide a complete understanding of the disease pathogenesis.

## Author Contributions

Conceptualization: R.A.P, L.P, M.N.R, I.B, M.B and M.A. Investigation: R.A.P, L.P, D.S, C.Schultheiβ, M.H.C and C.Santiskulvong. Resources: L.Y, H.C, M.L, J.V.E and A.F. Data Analysis: R.A.P, L.P, M.N.R, M.H.C, D.S, C. Schultheiβ, I.B, M.B and M.A. Writing: R.A.P, L.P, M.N.R, M.H.C, I.B, M.B, M.A.

## Acknowledgements

We gratefully acknowledge support from the NIH awards P41 GM103712 (to IB) and R01 AI072726 and 3RO1AI072726-10S1 (to MA).

## Methods

### Patient Samples

Biobanked deidentified patient blood remnant samples as well as control samples were obtained from Massachusetts General Hospital (Boston), Cedars Sinai Medical Center (Los Angeles), Loma Linda University Hospital (Loma Linda) and Martin-Luther-University Halle-Wittenberg, (Halle, Saale; Germany) under Ethics committee approval and informed consent. Samples consisted of mild MIS-C (n=4), severe MIS-C (n=16) and age-matched febrile controls patients (n=15). MIS-C severity classification was based on admission to Pediatric ICU requiring vasopressor use (severe) or no ICU admission (mild). Median age of febrile controls, mild and severe MISC were 12, 9 and 10.5 respectively. 95% of patients were serology positive for SARS-CoV-2 and 30% of patients were PCR positive by nasopharyngeal swab for SARS-CoV-2.

### TCR immunosequencing

RNA was isolated form peripheral blood of febrile controls and MIS-C patients and assessed for quality with a bioanalyzer (Agilent). Sequencing of the TCR genes was performed using the QIAseq Immune Repertoire RNA Library Kit (Qiagen) and the NovaSeq6000 system (2 x 250bp, 11M average reads per sample).

In an additional cohort of 254 healthy individuals of all age groups, the TRB genetic locus was amplified using 500 ng DNA from peripheral blood mononuclear cells (PBMC) in two consecutive PCR reactions using BIOMED2 TRB A/B primer pools (van Dongen et al., 2003). During the first PCR, fragments were tagged with Illumina-compatible adapters for hybridization to the flow cell. After bead purification (AMPure XP, Beckmann Coulter, CA, USA) each sample was tagged with a unique 7 nucleotide index during the second PCR. All PCRs were performed using Phusion HS II (Thermo Fisher Scientific Inc., Darmstadt, Germany). After the second PCR, amplicons were separated by gel electrophoresis and purified using NucleoSpin^®^ Gel and PCR Clean-up (Macherey-Nagel, Düren, Germany). Library quantification and quality control was performed using Qubit (QIAGEN, Hilden, Germany) and Agilent 2100 Bioanalyzer (Agilent Technologies, Böblingen, Germany), respectively.

### TCR immunosequencing analysis

The MiXCR framework (3.0.8) (Bolotin et al., 2015) was used for annotation of TCR rearrangements and clone construction, whereby the default MiXCR library served as reference for sequence alignment and each unique complementarity-determining region 3 (CDR3) nucleotide sequence was defined as one clone. Only productive sequences with a read count of ?2 were considered for further analysis. To account for differences in sequencing depth, TRB and TRA repertoires were normalized to 1.5 million reads; TRG and TRD were normalized to 50,000 reads per repertoire. All analyses and data plotting were performed using R version 3.5.1. Broad repertoire metrics (clonality, diversity, richness), gene usage and repertoire overlap were analyzed using packages tcR (Nazarov et al., 2015) and ade4 (Dray and Dufour, 2007) as previously described (Simnica et al., 2019b; Simnica et al., 2019c). CDR3 length distributions were visualized with package pheatmap (Kolde, 2019) and represent the clonal abundance within each repertoire.

Bubble diagrams, which depict the clonal distribution of repertoires, were generated using packages packcircles (Bedward et al., 2018) and ggplot2 (Wickham, 2016). For plotting purposes, repertoires were sampled down to 20,000 reads. The area size of the bubbles reflects the clonal fraction.

CDR3 amino acid sequence similarity was assessed using Gapped local alignment of Motifs (GLAM2, (Frith et al., 2008)) and FastTreeMP (Price et al., 2010), respectively. For GLAM2 the 50 most abundant CDR3 amino acid sequences of TRBV11-2 clones were used as input to generate the alignment LOGOs. Another alignment approach was to focus on the most frequent 100 clones of each repertoire, irrespective of V gene. The amino acid sequence covering framework region (FR) 2 to CDR3 or CDR3 only of the rearranged TRB locus was used to infer sequence similarities with FastTreeMP (Price et al., 2010) and the data were visualized and analyzed using the Archaeopteryx viewer (0.9928 beta) (Han and Zmasek, 2009).

### Serum Cytokine Analysis

Quantification of cytokines in serum was performed with a highly sensitive multiplex enzyme-linked immunosorbent assay (ELISA) using electrochemiluminescence detection technology (Meso Scale Discovery [MSD], Rockville, Maryland, USA). Cytokine correlation with TRBV11-2 usage was assessed by Spearman r correlation test.

### In silico structural modeling

The structural model for SARS-CoV-2 spike was generated using SwissModel(Waterhouse et al., 2018) based on the resolved cryo-EM structures (Protein Data Bank (PDBs): 6VSB(Wrapp et al., 2020) and 6VXX(Walls et al., 2020)). The structure of the T cell receptor (TCR) with the Vβ domain 100% sequence identity to TRBV11-2 was taken from the crystal structure of HLA B*0801 in complex with a peptide (HSKKKCDEL) and DD31 TCR (PDB: 4QRP)(Nivarthi et al., 2014). Both a- and β-chains of the TCR(Nivarthi et al., 2014) (respective chains D and E in PDB 4QRP) were adopted for docking.

Generation of the TCR-spike binary and TCR-MHCII-spike ternary complexes: Using protein-protein docking software ClusPro (Kozakov et al., 2017), we constructed *in silico* a series of binary complexes for SARS-CoV-2 spike and TCR-TRBV11-2, following the protocol we used in our previous study (Cheng et al., 2020). Given that the constant domain is proximal to the cell membrane and TCR employs the variable domain for binding superantigens (SAgs) and/or antigen/MHC complexes (Saline et al., 2010), we added restraints to our docking simulations to prevent the binding of the TCR constant domain. An ensemble of 460 models was generated for the complex, which were grouped in 30 clusters based on their structural similarities. Among them, three displayed binding between the spike and TRBV11-2 at the SAg-like region containing the polybasic insert PRRA. The TCR-MHCII-spike ternary complexes were generated following the protocol described in previous work (Cheng et al., 2020). In brief, the 3-dimensional structure of MHCII was taken from the crystal structure of the ternary complex (Saline et al., 2010) (PDB: 2XN9) between human TCR, *staphylococcal* enterotoxin H (SEH), and MHCII. First, we performed docking simulations to generate binary complexes between MHCII and SARS-CoV-2 spike. Six representative MHCII-spike binary complexes were selected to explore binding of TCR to form a ternary complex. Potential ternary MHCII-spike-TCR complex models were selected from a generated ensemble of conformers following three filtering criteria: (i) TCR binds to the “PRRA” insert region; (ii) the binding region of spike preferably includes one or more segments that are sequentially homologous to the SAg-like or toxin-binding motifs predicted for SARS-CoV-2; (iii) MHCII and TCR are in close proximity. Figure 5C displays a representative conformer compatible with the binary complex shown in Figure 5A.

