## Supplemental Figures for "Identification of a unique TCR repertoire, consistent with a superantigen selection process in Children with Multi-system Inflammatory Syndrome"

Supplemental Figure 1

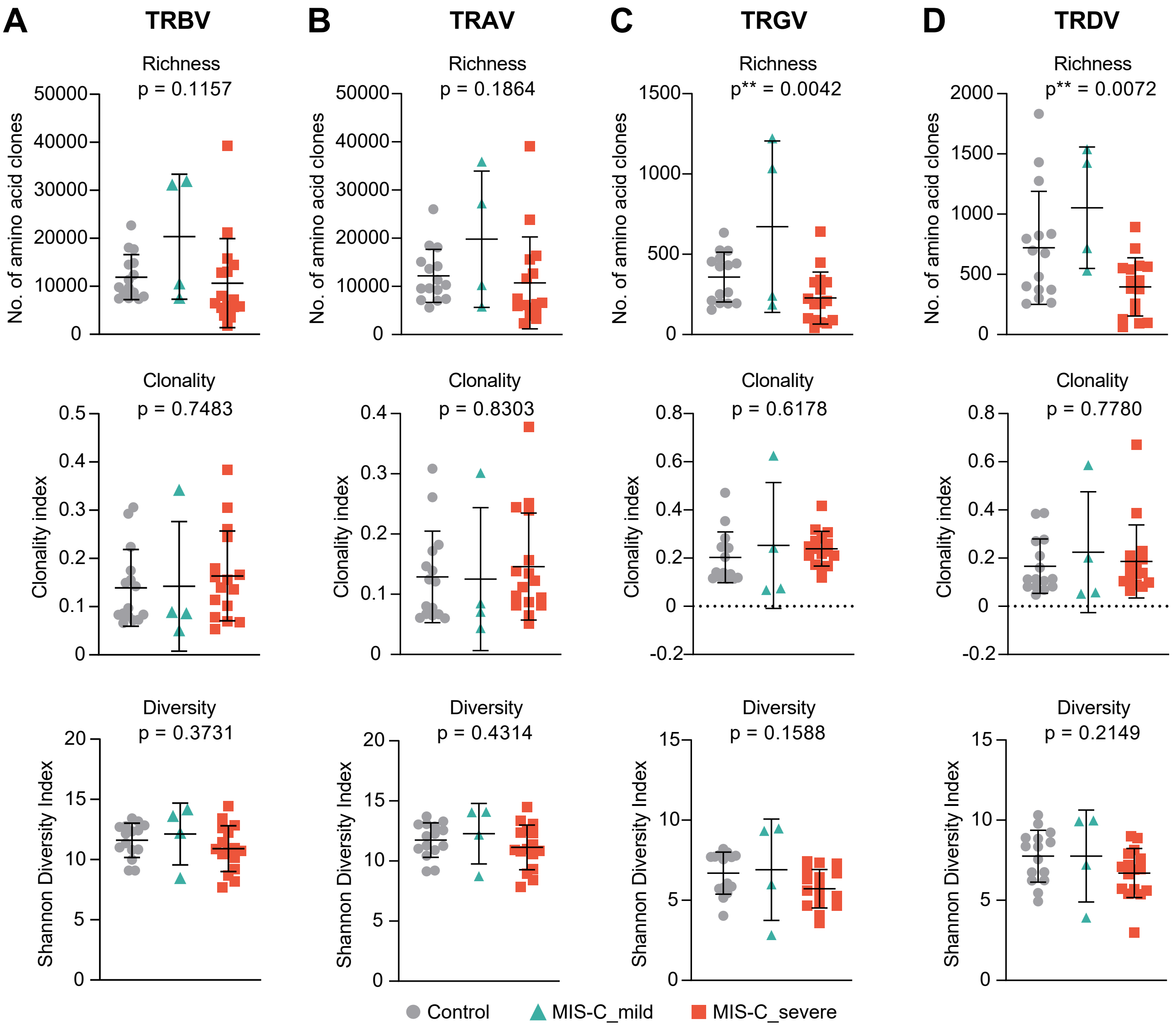


**Figure S1. T Cell Repertoire Metrics of MIS-C patients, Related to Figure 1.** Richness, clonality and Shannon diversity of the productive TRBV, TRAV, TRGV and TRDV repertoires derived from the peripheral blood of mild (n=4) and severe MIS-C (n=16) patients were compared to the respective T cell repertoires derived from the blood of age-matched febrile control patients (n=15). Bars indicate mean ± SD. Statistical analysis: Ordinary one-way ANOVA.

Supplemental Figure 2

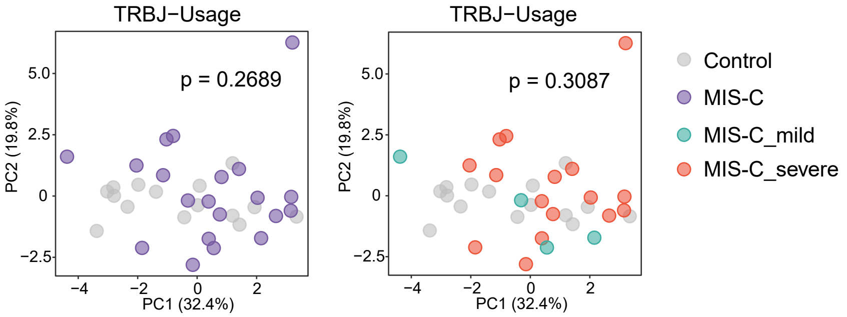


**Figure S2. Principal Component Analysis of TRBJ Usage, Related to Figure 1.** TRBJ usage in MIS-C patients (n=20) versus febrile controls (n=15) and between mild MIS-C (n=4) and severe MIS-C (n=16) patients. Statistical analysis: Pillai–Bartlett test of MANOVA of all principal components.

Supplemental Figure 3

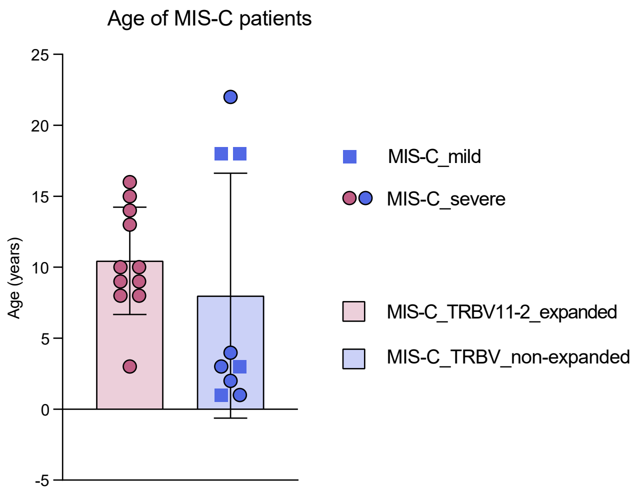


**Figure S3. Association of Patient Age with MIS-C Severity and TRBV11-2 Expansions, Related to Figure 2.**

### Supplemental Figure 4

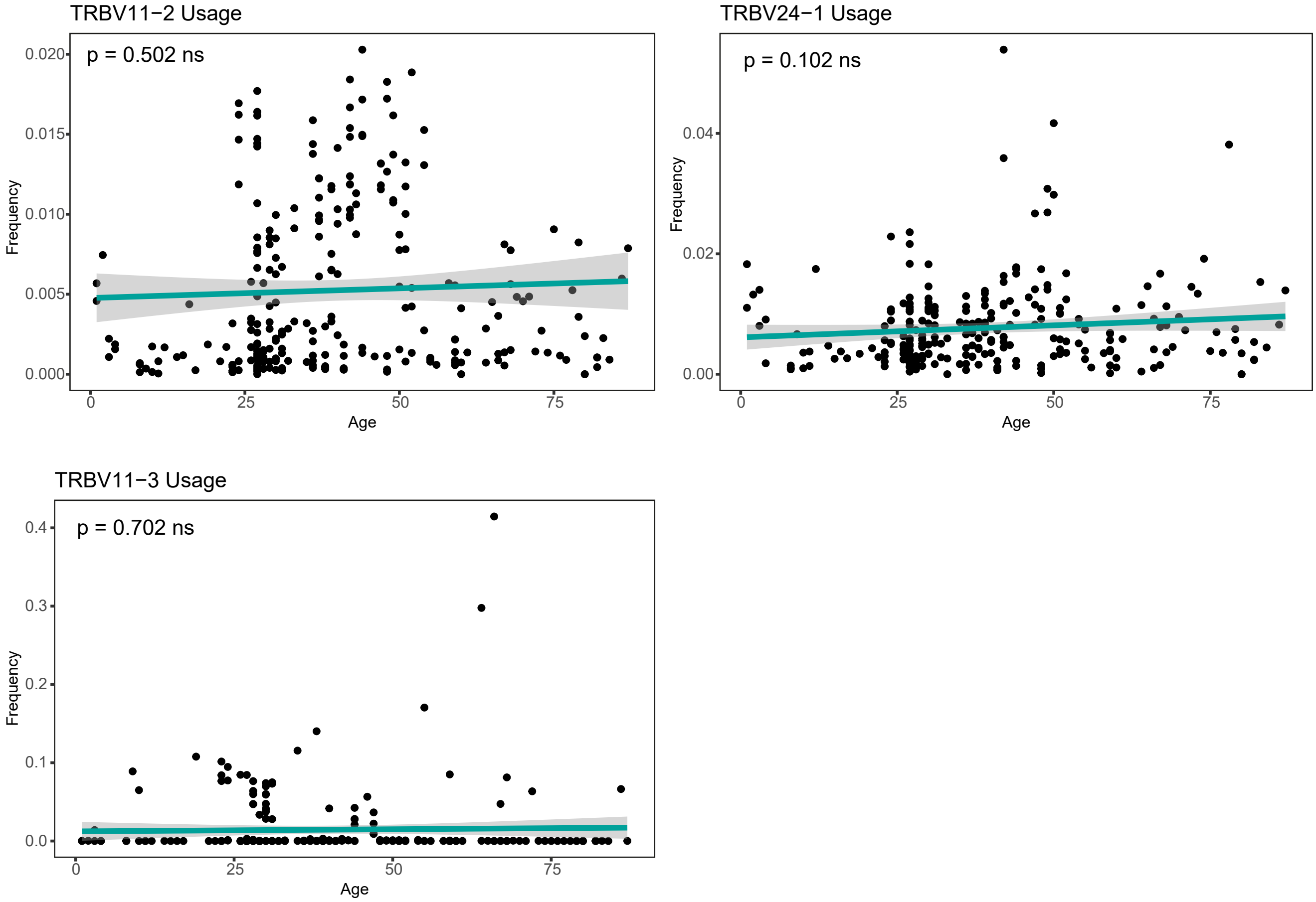


**Figure S4. Association of TRBV11-2 and TRBV24-1 Gene Usage, Related to Figure 2.** Frequencies of rearranged TRBV11-2, TRBV24-1 and TRBV11-3 genes in 254 repertoires of healthy individuals of all age groups plotted against age. Statistical analysis: Pearson correlation.

Supplemental Figure 5

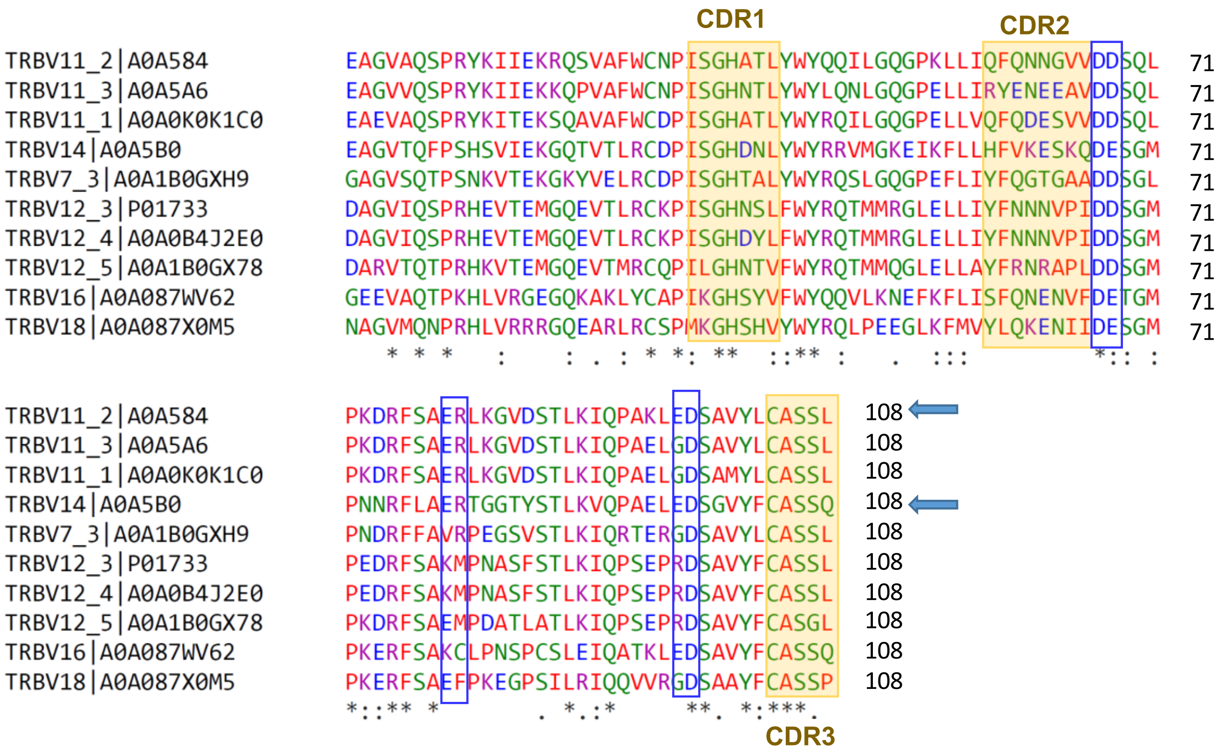


**Figure S5: Multiple Sequence Alignment of all TRC Vβ Chains that Contain Polyacidic Residue Pairs D67-D68 or D67-E68 Near their CDR2 Region, Related to Figure 5.** TRC Vβ sequences containing polyacidic residues at their CDRs were retrieved from the IMGT Repertoire. The alignment was generated using Clustal Omega (Sievers et al., 2011). CDR1, CDR2 and CDR3 are indicated by *orange shades*.
